# Molecular analyses reveal consistent food web structure with elevation in rainforest *Drosophila* - parasitoid communities

**DOI:** 10.1101/2020.07.21.213678

**Authors:** Christopher T. Jeffs, J. Christopher D. Terry, Megan Higgie, Anna Jandová, Hana Konvičková, Joel J. Brown, Chia Hua Lue, Michele Schiffer, Eleanor K. O’Brien, Jon Bridle, Jan Hrcek, Owen T. Lewis

## Abstract

The analysis of interaction networks across spatial environmental gradients is a powerful approach to investigate the responses of communities to global change. Using a combination of DNA metabarcoding and traditional molecular methods we built bipartite *Drosophila*-parasitoid food webs from six Australian rainforest sites across gradients spanning 850 m in elevation and 5° Celsius in mean temperature. Our cost-effective hierarchical approach to network reconstruction separated the determination of host frequencies from the detection and quantification of interactions. The food webs comprised 5-9 host and 5-11 parasitoid species at each site, and showed a lower incidence of parasitism at high elevation. Despite considerable turnover in the relative abundance of host *Drosophila* species, and contrary to some previous results, we did not detect significant changes to fundamental metrics of network structure including nestedness and specialisation with elevation. Advances in community ecology depend on data from a combination of methodological approaches. It is therefore especially valuable to develop model study systems for sets of closely-interacting species that are diverse enough to be representative, yet still amenable to field and laboratory experiments.

## Introduction

The geographic distribution of a species is determined by its intrinsic abiotic limits and by its interactions with other species. Individual populations of species do not respond to environmental changes in isolation from the species with which they interact (Davis et al. 1998, Gilman et al. 2010, Urban et al. 2016). Furthermore, interspecific interactions themselves respond to environmental change in diverse ways (Tylianakis et al. 2008, Thierry et al. 2019, Guiden et al. 2019). To understand how species’ distributions will respond to global change, it is therefore important to consider this wider community context.

One valuable approach for understanding how ecological communities will respond to environmental change is to study how food webs and other networks of interspecific interactions vary along elevational gradients (Tylianakis and Morris 2017, Pellissier et al. 2018, Baiser et al. 2019, Gravel et al. 2019). Temperature changes rapidly with elevation, so transects spanning elevations on mountain slopes can approximate aspects of climate change, while avoiding many of the confounding factors such as dispersal barriers or differences in seasonality that complicate analyses of latitudinal gradients (Hodkinson 2005, Körner 2007, Malhi et al. 2010, Morris et al. 2015). There is an expectation that predation and parasitism risk increase with temperature (Roslin et al. 2017, Libra et al. 2019), but it is unclear how such changes are expressed at the community level. Quantitative food webs should be particularly valuable in exploring changes in species interactions with changes in temperature, because they incorporate the relative abundances and interaction frequencies of component species (Memmott and Godfray 1994). The structure of communities can be summarised with network-level metrics that can reveal shifts that are not apparent in simple connectance food webs or in qualitative descriptors of diversity such as species richness (Tylianakis et al. 2007, Delmas et al. 2019).

To date, there have been relatively few network analyses along elevational transects (Pellissier et al. 2018), in part because of the substantial research effort required to document them through direct observation (for example, by observing herbivores feeding on plants, or parasites associated with hosts). These traditional approaches to describing networks are highly labour-intensive, even for a single network. Rapid, accurate and cost-effective methods to characterise interaction networks are therefore needed to assess the community-wide consequences of environmental change.

Recent developments in molecular methods offer the potential to document interaction networks with lower ‘per-interaction’ effort, while also addressing biases and deficiencies such as those arising from insufficient sampling. Traditional methods of food web construction are also often prone to sampling biases (Gibson et al. 2011) and can miss cryptic species and associated interactions (Derocles et al. 2015). By contrast, DNA-based approaches can be faster, more efficient and taxonomically more comprehensive, allowing the simultaneous resolution of interactions and identification of morphologically-cryptic species (Hrček and Godfray 2015). Molecular approaches to the construction of ecological networks have become increasingly common over the last decade (Clare 2014, Evans et al. 2016, Derocles et al. 2018, Roslin et al. 2019, Evans and Kitson 2020). Although high throughput, ‘next generation sequencing’ (NGS) methods are now routinely used to characterise bulk samples, their use for characterising trophic interactions in food webs has proved more challenging. Recently-developed nested DNA metabarcoding allows large numbers of pooled samples to be processed in a single NGS run without losing resolution to individual samples (Kitson et al. 2019). This method has been used successfully to identify interactions involving a single species and its consumers (Kitson et al. 2019). However, it is unclear whether such success is scalable to more complex interaction networks.

Food webs involving insect parasitoids, whose larvae feed exclusively on, and kill, a single arthropod host (Godfray 1994), have considerable potential for studying the processes that structure communities as they are both experimentally tractable while retaining meaningful levels of complexity (Hrček and Godfray 2015). Host-parasitoid interactions are amenable to molecular identification (Hrček et al. 2011, Wirta et al. 2014, Derocles et al. 2015, Kitson et al. 2019), with the caveat that the method describes parasitoid attack rate rather than the rate of successful parasitism *per se*.

Host-parasitoid interactions are important drivers of insect population dynamics (Hassell 2000) and climate change is expected to cause distribution shifts in parasitoids (Burrows et al. 2014), alter host-location and utilisation, and change the balance between parasitoid virulence and the immune response of their hosts (Hance et al. 2007). The consequences for host-parasitoid communities and the functions and services that they mediate are likely to be pronounced (Jeffs and Lewis 2013, Derocles et al. 2018, Thierry et al. 2019). Previous work has found increased specialisation within host-parasitoid networks with increasing elevation (Maunsell et al. 2015). However, other sources of information about the effect of temperature, such as latitudinal trends, do not show a consistent trend in specialisation or other community-level properties (Morris et al. 2014, Moles and Ollerton 2016). Meta-analyses have found scale dependence of patterns (Galiana et al. 2019), but there is a need for additional studies to assess general trends in network structure.

Here we explore how food webs might respond to predicted climate change by describing the community of wild frugivorous vinegar flies (Diptera: Drosophilidae) and their parasitoids across two elevational transects in Australian rainforest. Biotic interactions (O’Brien et al. 2017, 2020) and thermal tolerance (Kellermann et al. 2012) have been shown to determine *Drosophila* species ranges. In this study, we resolve and quantify food web composition and structure using molecular methods. We demonstrate how nested DNA metabarcoding can be used to generate multiple quantitative networks across landscapes, opening up opportunities to use this and similar systems as models in community ecology research.

## Methods

### Study sites

The Australian Wet Tropics World Heritage Area is an area of rainforest with exceptional levels of biodiversity (Stork et al. 2011) which lies close to Queensland’s northeast coast between Cooktown and Townsville (15-19’S, 145-146.30’E). We established study sites along two transects through the rainforest along elevation gradients at Paluma Range Road in Paluma Range National Park and Kirrama Range Road within Girramay National Park (Figure 1). Our study sites spanned elevations from 59 – 916 m above sea level and mean temperatures ranged from 21°C at the higher elevation sites to 26 °C at lower sites (SI 1). Variability in temperature was not strongly related to elevation over this period, although the lower elevations show higher variation annually. The temperature differences across our elevation transects are broadly consistent with temperature shifts expected over the next 50-100 years under models of climate change (IPCC 2014). We selected seven forested sites along each transect spanning the three ‘ecologically significant climatic zones’ identified by Webb and Tracey (1981) on the basis of changes in forest structure with elevation.

**Figure 1.**
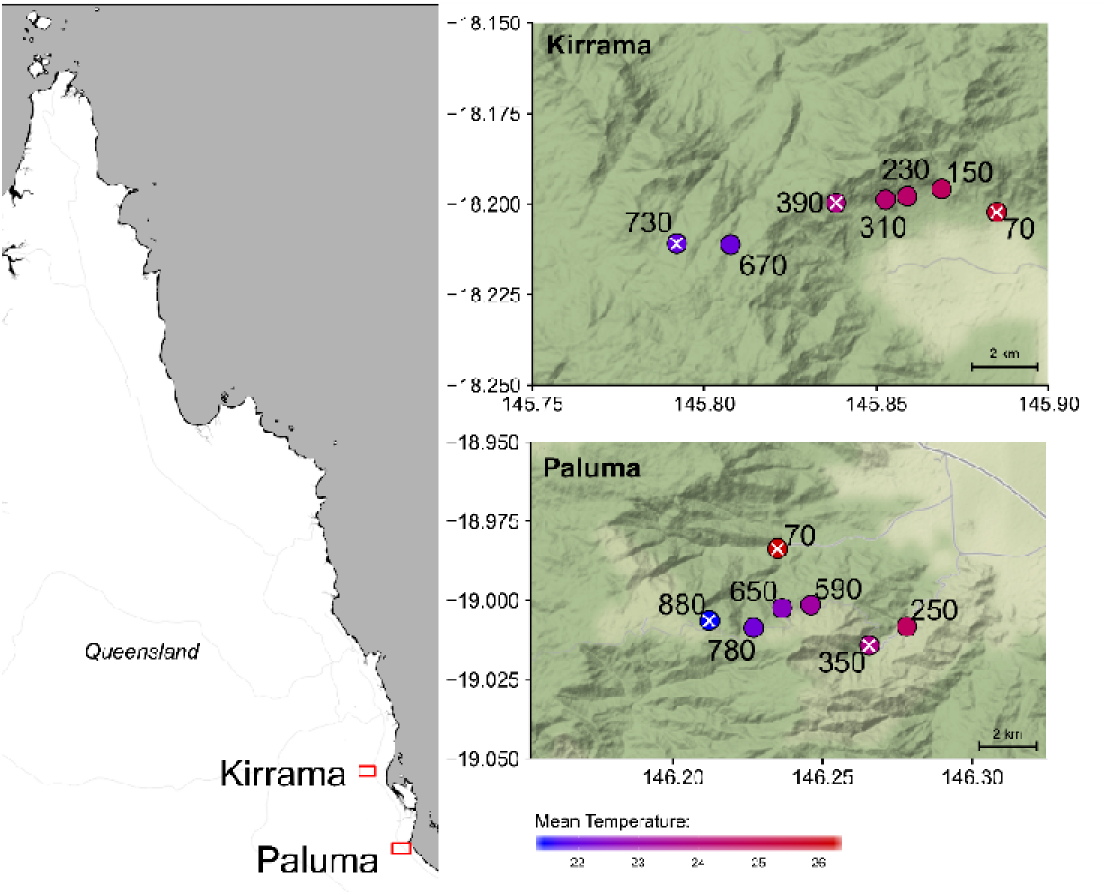
Location of transects in north-eastern Australia. Location of sites are coloured by the mean temperature recorded by dataloggers at each site during the research period, and labelled by the elevation (metres above sea level). White crosses show sites for which we constructed interaction networks. Map tiles by Stamen Design, under CC BY 3.0. Data by OpenStreetMap, under ODbL.

### Study system

We focused on the food web of interactions involving native, forest-dwelling, fruit-feeding *Drosophila* species and their associated parasitoids (Hymenoptera). Well-established laboratory practices and fundamental knowledge of *Drosophila* and their parasitoids (Kraaijeveld and Godfray 1997, Markow and O’Grady 2008, Prevost 2009) make these interacting species a valuable model system in community ecology. In our study area, *Drosophila* assemblages associated with rotting fruit are well characterised (Hangartner et al. 2015), although less is known about their parasitoids.

### Sampling Method

At each site, *Drosophila*-parasitoid communities were sampled using bottle traps baited with fermented banana. Bottle traps consisted of 1.5 L plastic water bottles with two 8cm × 10cm windows cut in the side to allow flies and wasps to enter, following Markow and O’Grady (2006). A rain-shield was placed over the lid of the bottle and the line was coated with Tanglefoot^®^ insect barrier (The Tanglefoot Company, Grand Rapids, USA) to prevent ants compromising the traps (see image in SI 1). Bottle traps were hung 1.5 m above the ground and separated by a minimum of 5 m. We set the traps in shaded locations at least 5 m from a roadside edge or canopy gap.

Previous work has established that banana bait samples the native generalist *Drosophila* spp. in our focal communities effectively (*Schiffer pers. obs*., Carson and Okada (1983). Traps were baited with 50 g of banana bait (mashed ‘Cavendish’ bananas supplemented with bakers’ yeast), and strips of cardboard were added to facilitate *Drosophila* pupation. Traps were exposed during the period 11^th^ March – 12^th^ April 2016, corresponding to mid-late wet season when insect activity is high. Parasitoids of *Drosophila* attack either larvae or pupae, but all emerge from pupae. Thus, by sampling *Drosophila* pupae we were able to document the network of host-parasitoid trophic interactions because pupae contain information about both the host and any parasitoid infecting it.

To capture the variation in colonisation speed and pupation time of different *Drosophila* species and across the elevation gradient, traps were open for either long (24 days), medium (14-15 days), or short (11-12 days) exposure periods at all sites. We did not find evidence of exhaustion of baits, indicating that competition for food was unlikely to consistently exclude components of the community. The relative frequency of the host species likely changes through the stages of the lifecycle through the differential impact of competition and lifespan. Our surplus-food design therefore most closely represents the relative frequency of the natural population in the early larval stage, before larval competition acts.

For each of the sites we sorted fly pupae from 15 randomly selected trap samples, five from each of the three exposure-time categories. We excluded traps if they had been compromised by ants or mammals, were exceptionally dry, or had been flooded by rain. Traps were either sorted on the day of collection or frozen at −15°C and sorted later. Pupae were stored individually in separate wells of 96-well plates filled with 96% ethanol and stored at −15°C. A random subset of pupae from the high and low elevation sites were reared in sealed, ventilated 96-well plates in a 22°C incubator to yield adult fly and parasitoid material. These were identified morphologically (*Drosophila* to species level by MS and parasitoid to genus level by C-HL) and used to build a DNA reference library (Table S2.1).

### Hierarchical Network Construction Approach

Quantitative interaction networks were built for three sites from both transects representing the highest, lowest and most central elevations. Pupa samples were randomly drawn from across the three exposure times in proportion to the number caught in each. Quantitative networks were generated by a hierarchical sampling strategy. First, a ‘core’ network was built from a random selection of 182 pupae per site, where both host and parasitoid (if present) were identified by molecular methods (detailed below). In this stage of network construction, 722 hosts were successfully identified in 86-138 samples per site. Of these, a total of 179 were found to be parasitoid positive using PCR, and in 147 cases the parasitoids were successfully identified.

Next, to optimise the number of interactions sampled from each network, all available additional samples (total 2659, 339-564 per site) were screened for parasitoid DNA using PCR. Those found to be parasitized were then sequenced to identify the host and parasitoid. This resulted in an additional 289 identified interactions (15-119 per network). These two sets of data were combined into an overall ‘enriched’ quantitative network (Figure S2.2). This used interaction frequencies derived from both sets of data, scaled by relative host frequencies and overall host-specific parasitism rates from the ‘core’ network data. In 5 cases an observed interaction in the enriching set could not be associated with a parasitism rate as that host was not observed to be parasitised in the core dataset at that site. These interactions were assigned a relative frequency of 0 in the enriched network. However, these interactions were included in the specification of the site-specific binary interaction matrices used for the calculation of site dissimilarity and qualitative network metrics.

### Molecular methods to describe network structure

We used a combination of DNA metabarcoding and classic molecular methods. We first performed DNA metabarcoding on a selection of individually tagged pupal samples (“core” samples). By comparing the sequences to a custom reference library, we were able to identify hosts and estimate overall host diversity. Contrary to expectations based on previous work (Kitson et al. 2019), metabarcoding did not allow parasitoid identification and therefore we could not use this approach to document and quantify host-parasitoid interactions. We therefore developed a number of classic molecular tools to obtain this missing information and allow cost-effective identification of thousands of samples. The classic tools were: i) multiplex PCR to identify the host species, followed by ii) Sanger sequencing of samples not identified by the multiplex, and iii) parasitoid detection and identification using PCR and Sanger sequencing. Full details are given in Supplementary Information 2.

### Patterns in *Drosophila* occurrence and parasitism rate

To measure rates of parasitism, we supplemented the data described above with additional screening results from 350 samples from each of the remaining four sampling sites along each transect, generating a grand total of 6780 parasitism assays. To explore trends in parasitism we fit minimal generalised linear models (GLMs) with a quasi-binomial error distribution, using elevation (metres a.s.l.) and transect (Kirrama or Paluma) as predictors. The significance of individual terms was assessed using F-tests. Since host-species turnover may mask or drive underlying elevational effects if species have different susceptibilities, we also fitted the model including *Drosophila* species as an additional categorical predictor. This analysis was restricted to three abundant *Drosophila* species observed at all six core sites (*D. pallidifrons, D. pseudoananassae* and *D. rubida*).

### Network metrics across the elevation gradient

We calculated five standard network statistics for each site, using the *bipartite* R package (Dormann et al. 2008): *H2*’, which measures overall level of specialisation in the network (Blüthgen et al. 2006); *weighted NODF*, a metric of nestedness that corrects for network size and connectance (Almeida-Neto et al. 2008); *quantitative modularity*, which assesses the extent to which the network is divided into discrete compartments (Beckett 2016); *network vulnerability*, the mean number of parasitoid species attacking each host; and network generality, the mean number of host species that each parasitoid attacks. We tested for the impact of elevation on each metric with a linear model, including the matrix size (number of hosts × number of parasitoids, following Morris et al. (2014) and transect (Kirrama or Paluma) as additional predictors.

To assess the precision with which we can infer these metrics given our sample sizes, we used bootstrapping of our available observations. For the enriched networks, we resampled each sampling stage separately. Bootstrapping network in this manner provides an indication of the degree of uncertainty in a metric value. However, because the networks can only become smaller, the results are not unbiased and should be interpreted with caution.

### Drivers of interaction turnover

Interaction *β*-diversity represents differences in the set of observed interactions between sites (Poisot et al. 2012, Graham and Weinstein 2018). Whole-network dissimilarity (*β*_*WN*_) between two sites can be partitioned into that attributable to species not being present at both sites (*β*_*ST*_) and that attributable to interactions only occurring at one site despite both species being present at both (*β*_*OS*_). Because methods partitioning quantitative similarity measures are not yet well established, we used the qualitative Jaccard similarity metric (Novotny 2009) and a customised version of the betalinkr function from the bipartite R package that takes into account host species observed at a site that were not observed interacting with parasitoids. We enforced *β*_*WN*_ = *β*_*ST*_ + *β*_*OS*_ by dividing *β*_*ST*_ and *β*_*OS*_ through by *β*_*WN*_/(*β*_*ST*_ + *β*_*OS*_) following Poisot et al. (2012). For additional comparison, we calculated the Jaccard similarity between the host species found at each site (*β*_*Host*_). For each of these metrics, we calculated pairwise dissimilarities between each our six core sites. We used multiple regression on distance matrices (Legendre et al. 1994) via the ecodist R package (Goslee and Urban 2007) to investigate whether these dissimilarities are driven by elevational or between-transect differences.

## Results

### Parasitism Rate

Overall parasitism rate declined significantly with elevation across our 14 sites. For each 100 m rise in elevation, the probability of being parasitized decreased by 14% (coefficient estimate *β*_*elevation*_= −1.49 km^−1^, se = 0.49, Pr(>F_1,11_) = 0.00865). Parasitism was significantly higher overall at Paluma than at Kirrama (*β*_*Paluma*_ = 1.06, Pr(>F_1, 11_)= 0.0023). The elevational trend was largely driven by data from the Paluma transect (Figure 2a), although the interaction term between elevation and transect was not statistically significant (Pr(>F_1,10_) = 0.135). However, the high over-dispersion in the data (the dispersion parameter was estimated to be 14.8) makes it challenging to identify interaction terms with confidence.

**Figure 2.**
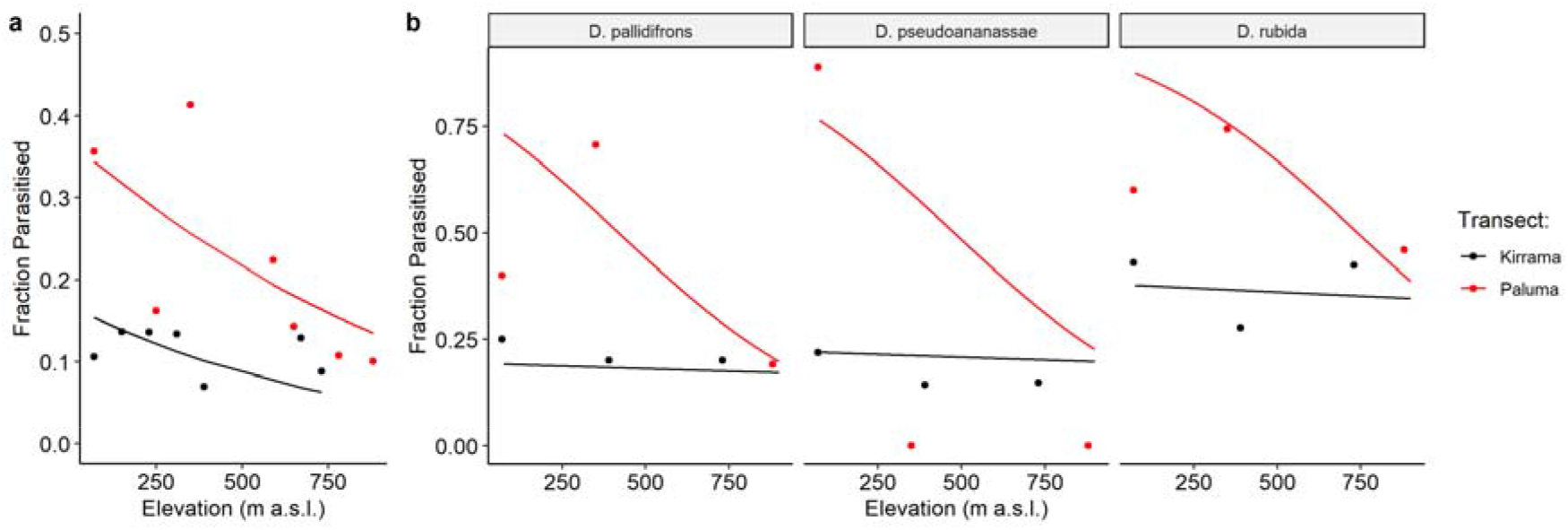
a) The fraction of pupae from traps at each site observed to be parasitized in relation to elevation. b) Parasitism rates at the six core sites for the three most abundant species in relation to elevation. In both subfigures, lines show the fitted minimal models.

Parasitism rates of widespread *Drosophila* species at the 6 focal sites showed similar patterns (Figure 2b). Model selection did not identify any host-specific transect (Pr(>F_2,8_) = 0.90) or elevation (Pr(>F_2,10_) =0.21) effects but did find a significantly greater decline in parasitism rate with elevation at Paluma (Pr(>F_1,11_) = 0.035). At Kirrama, there was only a negligible effect of elevation (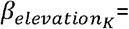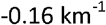, se: 0.85). Full statistical results and model selection procedures are given in SI 3.

### Network Structure

Across the six focal sites we found 12 *Drosophila* species, 15 parasitoid species (principally Figitidae, Diapriidae and Braconidae, Table S2.3), together making up 55 different interaction pairs. The host species found at Kirrama were a subset of those observed at Paluma. The six enriched quantitative networks are shown in Figure 3. There was no significant relationship between any of the tested network metrics and elevation (p>0.05 in all cases, Figure 4 & SI 3). Including network size as an additional co-predictor did not change this. Support intervals from resampling are shown in Figure 4.

**Figure 3.**
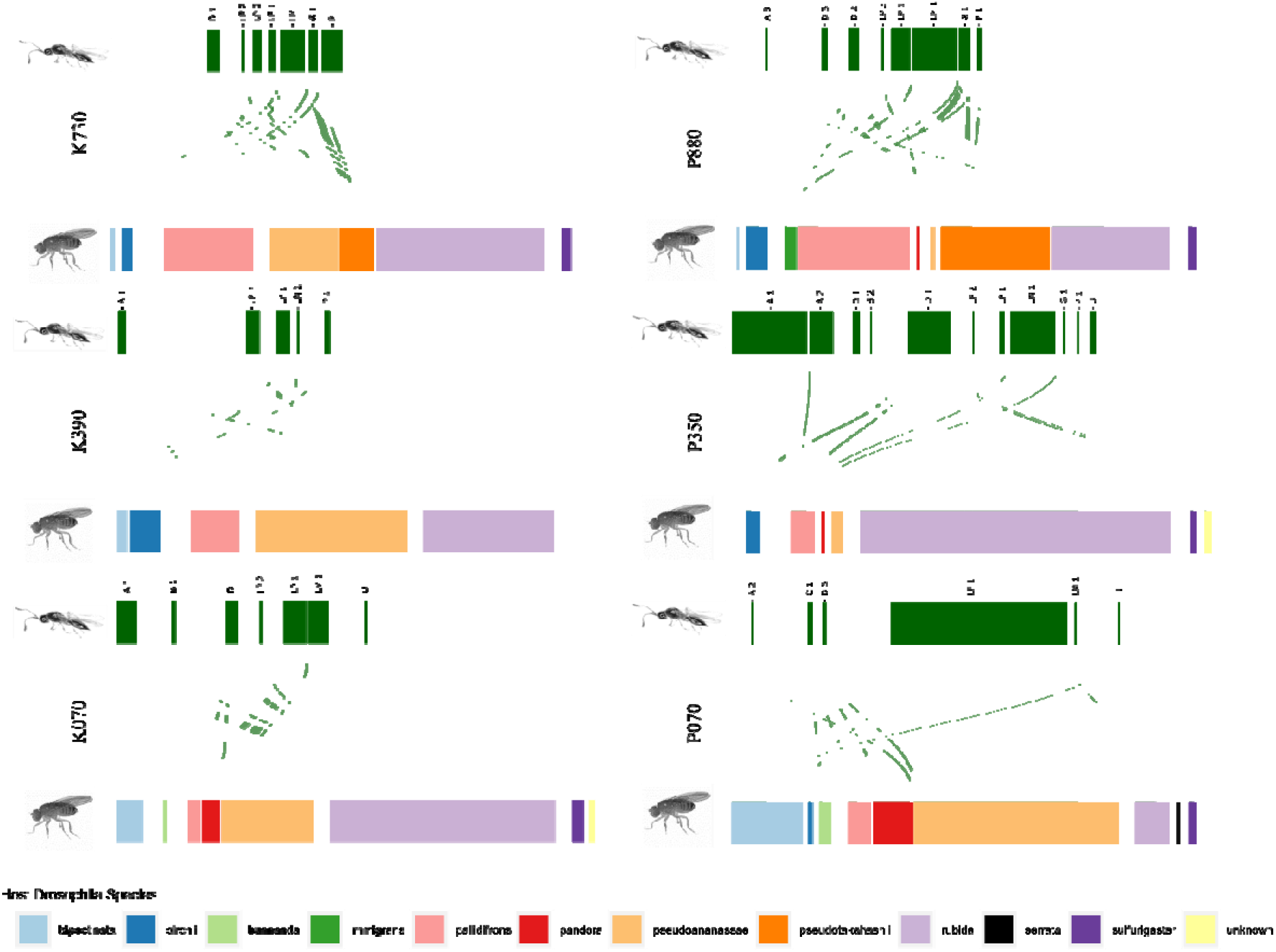
Quantitative interaction networks from each site, calculated using the enriching method described in the main text. The width of the boxes corresponds to the relative frequency that each species or interaction was observed. Networks are arranged in elevation order, with Kirrama on the left and Paluma on the right and labelled by the elevation of the site. Networks were drawn using the *bipartiteD3* R package. Parasitoid wasps are labelled by taxon: A= *Asobara*, B = Braconidae, C = Chalcidoidea, D= Diapriidae, G= *Ganapsis*, LM = *Leptolamina*, LP = *Leptopilina*, P = Pteromalidae, U = Unidentified.

**Figure 4.**
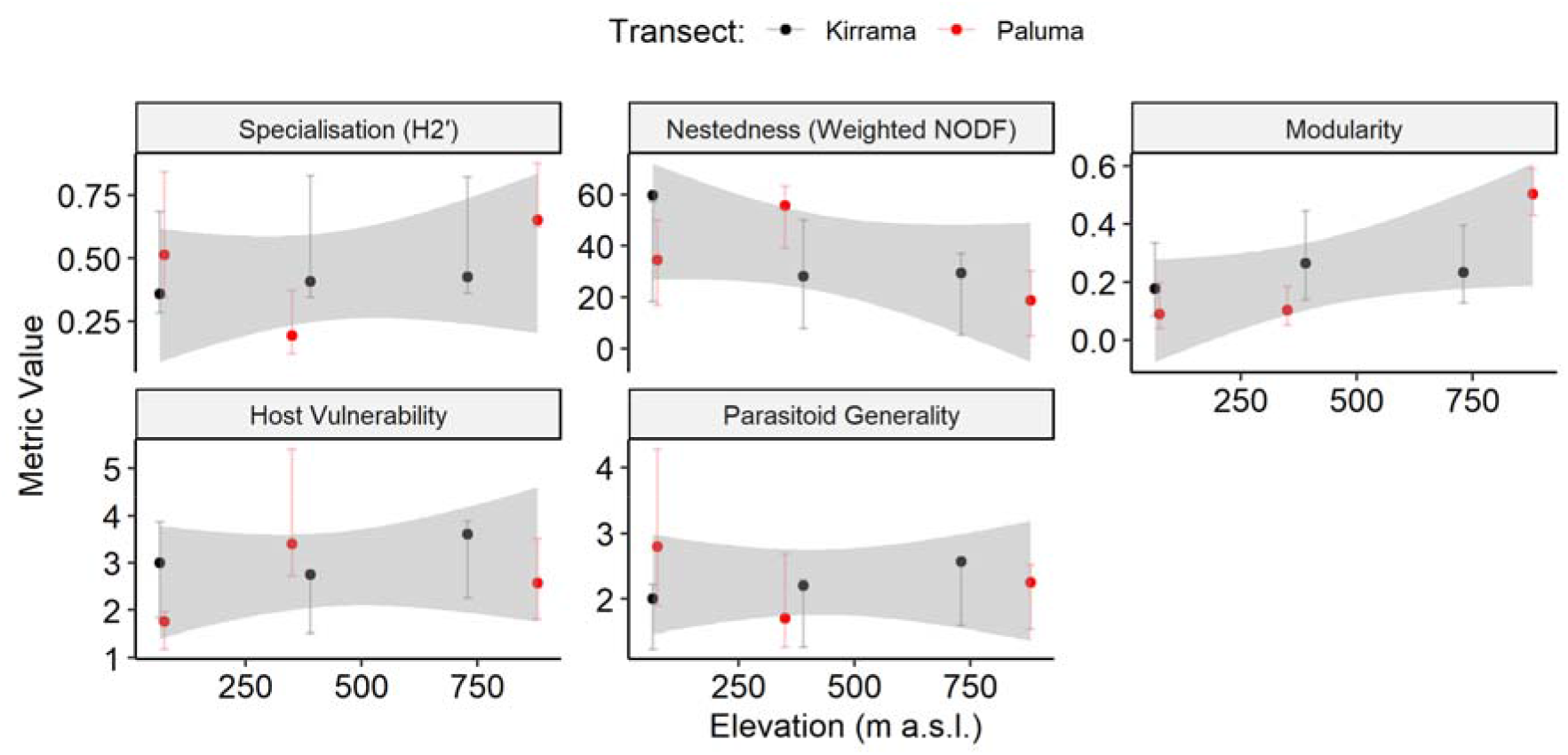
Consistency in network-level metrics across the elevational transect, with no statistically significant effect of elevation in all cases. Error bars show bootstrapped 95% support intervals from resampling the observations at each site. Shaded areas show 95% confidence intervals based on the elevation-only model.

### Drivers of network dissimilarity

Species turnover (*β*_*ST*_) was a more important driver of qualitative dissimilarity between networks (*β*_*WN*_) than changes in observed interactions (*β*_*OS*_, Figure 5a). Although dissimilarity tended to be higher for pairs of networks from locations differing greatly in elevation (Figure 5a), neither transect differences nor elevational differences significantly explained any of the components of network dissimilarity (all unadjusted p > 0.1). Overall host community composition disimilarity (*β*_*Host*_) was not significantly related to elevational differences (p = 0.45) or transect differences (p = 0.78).

**Figure 5.**
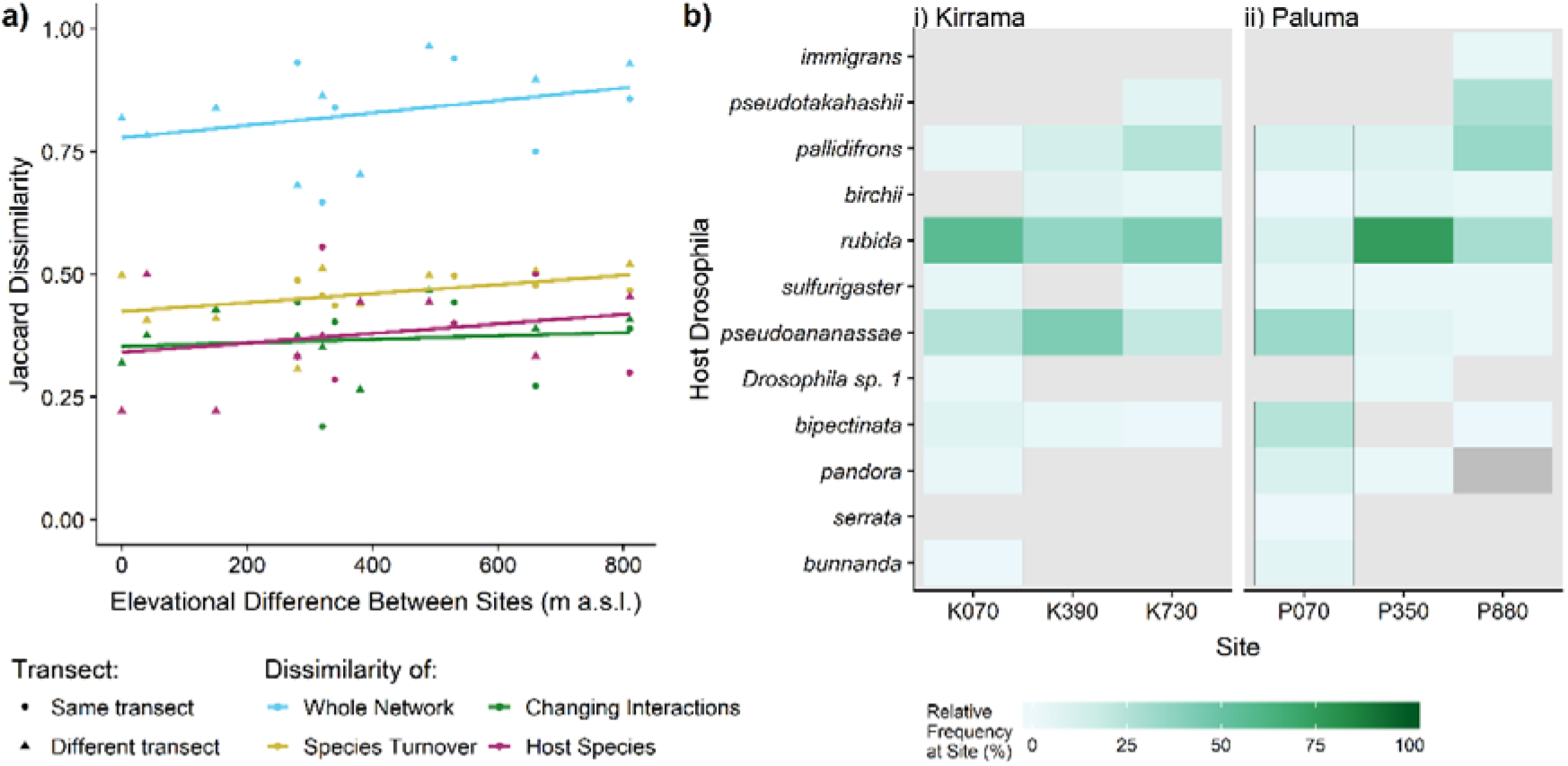
**a)** Effect of elevational difference on between-site community *β*-diversity measures. Whole-network interaction dissimilarity (*β*_*WN*_) was large, and predominantly composed of species turnover (*β*_*ST*_), although there was also a notable contribution derived from changing interactions between present species (*β*_*OS*_). Overall Jaccard dissimilarity in host species composition (*β*_*Host*_) was also notable. Although all the dissimilarity measures show an apparent increase with elevational difference, these trends were not significant. Lines show a direct linear model through each dissimilarity metric, ignoring transect differences. **b)** Relative observation frequency of the host species observed at each site. Host species are arranged in order of the centre of mass of their relative abundances. The dark grey square shows where *D. pandora* was observed in the enriching dataset, but not the core dataset, making its relative abundance indeterminable.

## Discussion

Our study demonstrates the capacity of molecular methods to sample multiple interaction networks of a model system efficiently and to shed light on the underlying ecological processes. We found evidence for a reduction in parasitism frequency at higher elevations in line with previous results from comparable systems (Kimura and Suwito 2015), although this was largely driven by one of our two transects. Despite this, we found no evidence for major changes to food web network structure along an elevation gradient that spanned a temperature range comparable to the ecological range of many of the focal species. This contrasts with the results from previous studies of host parasitoid food webs by Maunsell et al. (2015) and Morris et al. (2015) and suggests that any shifts in network structure with elevation are unlikely to be consistent and that, by extension, the response of host-parasitoid networks to climate change may not be easily predicted (Hance et al. 2007).

The lack of significant trends in network structure with elevation are consistent with the absence of marked trends in these metrics across latitudes documented by Morris et al. (2014) and the apparent constancy across landscapes in more focussed studies of antagonistic interaction networks (Kemp et al. 2017, Redmond et al. 2019).

Sampling networks across gradients involves unavoidable trade-offs when distributing sampling effort between sites, sampling periods and transects, in order to best capture any spatio-temporal changes in network structure. The sample size for each of our networks is still rather small, with a number of interactions represented by only a single observation. This makes it likely that many interactions remain unobserved (Jordano 2016). Nonetheless, the quantitative network metrics we used are frequently found to be amongst the most robust to moderate levels of sampling (**Banašek-Richter et al. 2004, Fründ et al. 2016, Vizentin-Bugoni et al. 2016, Henriksen et al. 2019**). With six sites, the power of our tests to detect small to moderate impacts of elevation is inevitably limited. Nevertheless, we did not detect a trend in any of the suite of five network structure metrics we tested. Any impact of elevation along our transect on network structure appears to be minor compared to that which we observed in the parasitism rate and species turnover. Identifying drivers of community change

We found a high degree of network dissimilarity between sites at both Kirrama and Paluma, which derived largely from species turnover (both host and parasitoid) rather than changes in the observed interactions. Although the transects have similar temperature profiles, the vegetation structure on the Kirrama transect had been more heavily disrupted by a cyclone five years before the sampling period. However, we did not identify any significant differences between the transects. We did not identify any significant effect of elevational differences on community differences, as measured by binary Jaccard dissimilarity. This pattern of higher-level consistency mirrors that found in a number of systems (Kaartinen and Roslin 2012, Kemp et al. 2017, Gravel et al. 2019, Redmond et al. 2019) and suggests that there are mechanisms channelling communities towards particular structures, despite external changes.

Although there are known shifts in relative abundance of host species in our study system with elevation (for example, *D. birchii* has higher abundance at cool, high-elevation sites than in warmer, low-elevation sites (O’Brien et al. 2017)), many of the host species occur across the full range of elevations sampled, leaving only a minority of high or low elevation specialists to drive any qualitative differences (Figure 5b). In terms of the interactions themselves, we found relatively high levels of generality – the mean level of specificity (as measured by *H2*’) in our communities (0.42) was notably lower than the mean of 0.65 found in a cross-latitude survey of host-parasitoid networks (Morris et al. 2014). Although the majority of species were found across the elevation range, their relative abundances varied considerably with elevation (Figure 5b). Differences in network composition may be difficult to detect with the binary network dissimilarity methods we used, and emphasise changes in comparatively rare species, which are more likely to be influenced by sampling effects (Chao et al. 2005). This highlights the importance of recording abundance rather than spatial range limits in determining species’ responses to temperature.

To address this challenge, Staniczenko et al. (2017) proposed that changes in relative interaction frequency could be used to link quantitative changes in interaction frequencies to environmental changes. We attempted to use such an approach within a Bayesian framework to relate elevation to per-capita interaction frequencies (*not shown*). However, we could not confidently identify any strong evidence for individual interactions being modified across the gradient, even when restricting analysis to the most widely observed interactions. Without independent external estimates of the relative abundance of parasitoid species, analysis of changing interaction rate is severely restricted. More fundamentally, variation in the identity and strength of links within networks along environmental gradients may be driven by any combination of turnover in species composition, changes in species abundances and by abiotic influences on species interactions (Pellissier et al. 2018). Moving from ‘how’ to ‘why’ is a critical challenge in network ecology, but feedbacks between the relative abundance of each species and the interactions between them generate fundamental circularities (Dormann et al. 2017, Moreira et al. 2018). An essential part of future work to identify the drivers of community composition will be controlled manipulative experiments (Derocles et al. 2018). This and other *Drosophila*-parasitoid systems offer exceptional potential for field manipulations (O’Brien et al. 2017, 2020), identification of physiological traits (Kellermann et al. 2012) and laboratory experiments (Davis et al. 1998).

### Sampling strategy and molecular methods

Despite the per-sample cost of molecular methods falling significantly in recent years, identification effort is still a finite and valuable resource. Our hierarchical sequencing approach was helpful in increasing the number of identified interactions in our dataset for a given amount of laboratory effort, an approach that can help to maximise scarce resources and particularly relevant in species-rich interaction networks. One drawback is that the additional data wrangling required to construct quantitative networks from the resulting data can complicate subsequent analyses, and in particular to propagate estimates of uncertainty in network structure.

Our sampling strategy was only a single snapshot of the interaction network at each of our sites, and therefore does not resolve seasonal or annual turnover in interactions (Kemp et al. 2017, Lue et al. 2018). Our approach of gathering pupae samples after a variety of exposure times enables us to characterise a community where development time varies among species and with temperature. However, a number of challenges remain. For example, shorter exposure periods are likely to underestimate the abundance of host species with longer development times. Species with rapid development times may therefore be over-represented in our sampling, although we did not detect consistent differences in the host composition between our exposure periods. Furthermore, rather than parasitism rate, our interactions are more accurately described as attack-rates, since it is not necessarily the case that a parasitoid egg or larva within a host would develop successfully, kill the host and emerge as an adult (Condon et al. 2014). Rearing to emergence is the only way to establish the result of a parasitism event (Hrček and Godfray 2015), but this is much more labour intensive, introduces other biases (such as survival in laboratory conditions), and would negate the advantages of the molecular approach.

DNA metabarcoding alone was unable to describe our host-parasitoid networks without supplementary use of conventional molecular methods. Although Kitson et al. (2019) demonstrated the DNA metabarcoding method on a single host and parasitoid species, we were unable to obtain network data with this method alone in our multispecies system. This was despite pure parasitoid DNA from our reference samples amplifying well with the primers we used. The reason is probably amplification bias to host DNA and low relative parasitoid/host DNA concentration. We estimate that increasing sequencing coverage per sample by several orders of magnitude would be necessary to characterise parasitoids and their links to hosts reliably. This would not currently be cost-effective, but might become more realistic in the near future as sequencing costs fall. We recommend using a single-step PCR with larger number of primer tags in future studies, since the nested metabarcoding approach involving two PCRs may be more prone to mistagging. We used a strategy for detecting mistagging developed by Kitson et al (2019) and found it to be a crucial aspect for producing reliable and verifiable DNA metabarcoding data. We therefore recommend incorporating mistagging controls to DNA metabarcoding runs in addition to negative and positive controls.

## Conclusion

We documented pronounced turnover of *Drosophila* species relative abundance across an elevation and temperature gradient. A strong reduction in parasitism with elevation was not associated with a detectable change in overall network structure. DNA metabarcoding proved useful to characterise these previously unstudied ecological networks, but it did not prove possible to use this approach “out of the box” to obtain network data. Rather, metabarcoding data had to be supplemented with data from classic molecular methods and a reliable DNA reference library. Ecological communities are highly complex and communities will have to respond to multiple changes under global change. To understand the patterns in these networks and to distinguish the principal drivers behind them, manipulative experiments in the field and laboratory will be needed. A focus on tractable model systems will allow an in-depth investigation into fundamental ecological dynamics of natural systems.

## Author Contributions

CTJ: Led fieldwork, drafting manuscript, sampling design, performed molecular work. JCDT: Led statistical analysis and writing of manuscript. MH: Contributed to funding application, supervised sampling and fieldwork. AJ: Contributed to development of molecular methods, performed molecular work, HK: Contributed to development of molecular methods, performed molecular work. JJB: Performed molecular work. CHL: Parasitoid taxonomy, built parasitoid reference library, MS: *Drosophila* taxonomy. EO-B: Contributed to funding application, sampling design and fieldwork. JB: Conceived project, secured funding, contributed to sampling design. JH: Secured funding, led molecular work, writing. OTL: Conceived project, secured funding, study design, writing. All authors contributed to editing of the final manuscript.

CTJ and JCDT are joint-first authors. JH and OTL are co-last authors.

## Acknowledgments

We thank Adam Bajgar, Michaela Borovanská, Eva Chocholová for help in the molecular lab; Melanie Thierry and Nicholas Pardikes for help with live flies and parasitoids for the barcoding library, Ary Hoffman for COI barcodes; Adriana Uzqueda, Jaimie Hopkins and Leah Carr for help with field collection and sample sorting; and Lexie Edwards for help with logistics. Fieldwork was conducted under permit WITK16977516 to CTJ from Queensland’s Department of Environment and Heritage Protection.

## Code and Data availability

Code to replicate all analyses is available on Github at https://github.com/jcdterry/AusDrosTransect_CodeandData and if accepted will be archived on Zenodo. Raw data is also archived in a long-term data repository (EIDC https://doi.org/10.5285/85657c4c-54c9-4d02-a262-455ea1c38d95).

