## Supplementary material for "Molecular analyses reveal consistent food web structure with elevation in rainforest *Drosophila* - parasitoid communities": SI 1, 2 and 3

Supplementary Information 1 - Site Information

### Trap Design

At each site Drosophila-parasitoid communities were sampled using bottle traps baited with fermented banana. Bottle traps consisted of 1.5 L plastic water bottles with two 8cm x 10cm windows cut in the side to allow flies and wasps to enter. A rain-shield was placed over the lid of the bottle and the line was coated with tanglefoot to prevent ant raids. Bottle traps were hung 1.5 m above the ground and separated by a minimum of 5 m. We set the traps at least 5 m from a roadside edge to minimise effects of direct sun exposure. Traps were baited with 50 g of banana bait (mashed ‘Cavendish’ bananas supplemented with bakers’ yeast), and strips of cardboard were added to facilitate *Drosophila* pupation.

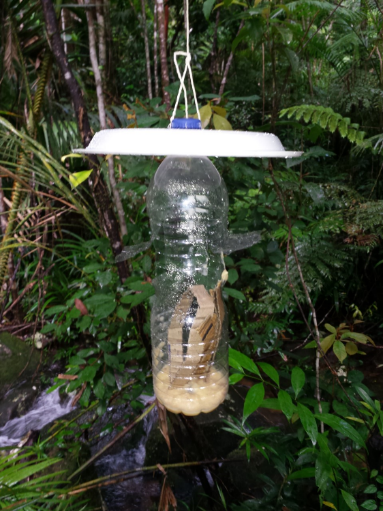

**Figure S1.1.** Traps used for pupae collection

The ‘medium exposure’ sampling period followed directly on from the ‘short-exposure’ sampling period, at the same locations. The ‘long-exposure’ sampling spanned the two briefer periods.

### Site Details

Three temperature and relative humidity dataloggers were attached to three randomly selected bait traps per site with a reading taken every hour for the duration of sampling. Data from dataloggers were used for Paluma between the period midnight 13th March - 11pm 1st April 2016, and data from Kirrama from between midnight 15th March – 11pm 3rd April 2016, the periods where all sites within each transect possessed full days in the field. Four Kirrama sites (K070, K150, K230, and K310) only had usable data from 18th March for two of the three dataloggers used. Paluma 070m was set up later than all other sites, and thus data presented are from 29th March - 11th April 2017. Changes in seasonal temperatures are not expected to alter the ranking of sites in relation to abiotic values, making elevation a good surrogate for climate in our analyses (Figures S1.2 and S1.3).

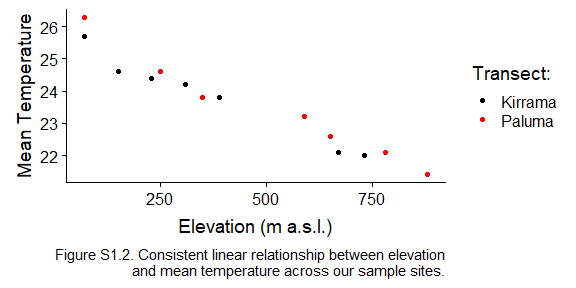

Table S1.1. Site location and mean temperature

| Site | Elevation (m) | Mean Temperature | Temperature standard deviation | Latitude | Longitude |
| --- | --- | --- | --- | --- | --- |
| K070 | 70 | 25.7 | 2.81 | -18.2022 | 145.8850 |
| K150 | 150 | 24.6 | 2.44 | -18.1959 | 145.8690 |
| K230 | 230 | 24.4 | 2.04 | -18.1979 | 145.8590 |
| K310 | 310 | 24.2 | 1.97 | -18.1988 | 145.8526 |
| K390 | 390 | 23.8 | 2.24 | -18.1997 | 145.8383 |
| K670 | 670 | 22.1 | 2.80 | -18.2112 | 145.8077 |
| K730 | 730 | 22.0 | 2.34 | -18.2109 | 145.7920 |
| P070 | 70 | 26.3 | 3.12 | -18.9837 | 146.2350 |
| P250 | 250 | 24.6 | 2.15 | -19.0083 | 146.2782 |
| P350 | 350 | 23.8 | 1.94 | -19.0143 | 146.2656 |
| P590 | 590 | 23.2 | 2.22 | -19.0015 | 146.2461 |
| P650 | 650 | 22.6 | 1.87 | -19.0025 | 146.2365 |
| P780 | 780 | 22.1 | 2.30 | -19.0087 | 146.2269 |
| P880 | 880 | 21.4 | 2.15 | -19.0064 | 146.2122 |

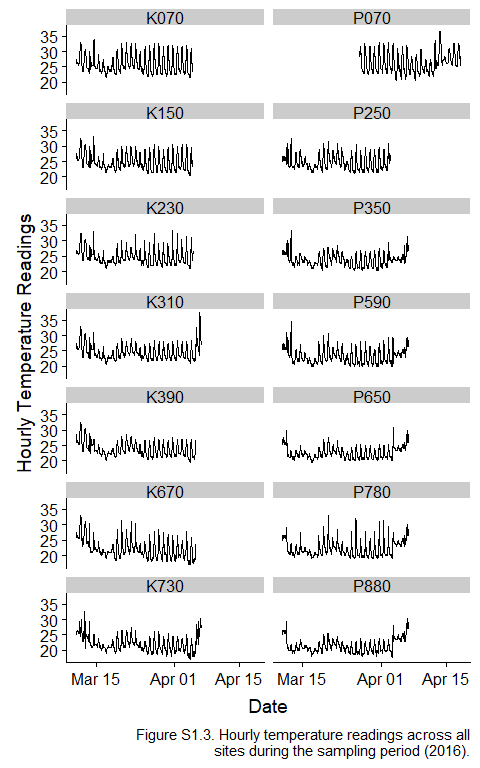

Supplementary Information 2 – Molecular Methods

**DNA Metabarcoding**

We largely followed the nested tagging DNA metabarcoding protocol developed by Kitson et al. (2019). DNA for metabarcoding was extracted from individual pupae using column extractions (GeneAid) in batches of 29 plus a negative control. We amplified a 421bp long segment of the COI barcode region using BF2-BR2 primers (Elbrecht and Leese 2017), modified by adding multiplexing indices as in Kitson et al (2019), including the use of separate indices for positive and negative controls which allows rigorous tracking of potential contamination and mistagging. We pooled 12 batches of 96 PCRs on 80% of one sequencing run. Each batch contained one positive control, two extraction negative controls, two PCR negative controls and 91 samples. The positive controls were extracted in the same way as normal samples and consisted of one *Drosophila* plus one wasp individual from stock lines of 12 *Drosophila* species and four wasp species collected at the focal sites and maintained in our laboratory. The samples, positive controls and negative controls were randomly placed within each batch and thus occupied different positions in different batches.

PCRs were carried out over 40 cycles (95°C for 15 s, 48°C for 15 s and 72°C for 30 s) in 15 µl reactions using a high fidelity Taq mastermix (MyFi Mix Bioline), 0.75 µl of template DNA and each primer (final concentration – 0.5 µM). Like Kitson et al (2019), we had to perform the protocol twice with additional measures preventing contamination of the PCR to obtain a clean sequencing run with good signal to noise ratio. We use data only from the second run. The additional measures were: 1) Single-use aliquots of forward and reverse primer mix for each batch were prepared in PCR box before the PCRs; 2) PCR was performed under a layer of mineral oil in PCR strips with single capped wells; 3) only single-channel pipettes were used and were decontaminated after each batch using 0.5% sodium hypochlorite solution. PCR results were checked on the gel. The differentially tagged PCR products within each batch were pooled in equal volume (8 µl) without DNA quantification, resulting into 12 pre-libraries corresponding to 12 batches which were submitted to Macrogen for sequencing. At Macrogen the pre-libraries were cleaned-up using AMPure XP beads, quality checked, and an additional tag was added using PCR with 8 cycles following Illumina Metagenomic Sequencing Library Preparation protocol (“Part # 15044223 Rev. B”). The sequencing library was then prepared from cleaned-up pre-libraries, equalizing the amount of PCR product per sample and sequenced on 80% of an Illumina MiSeq 300PE run with 5% PhiX.

Overall, the 80% of MiSeq run resulted in 14,838,676 reads. The data were quality trimmed to phred Q30 in Trimmomatic 0.32 (Bolger et al. 2014) using a sliding window approach (5 bp window size) and reads shorter than 100bp discarded. This step reduced the dataset to 11,495,448 reads (77%). Reads were then demultiplexed using STACKS (Catchen et al. 2013), and primers were clipped. Paired-end reads were subsequently merged (minimum overlap 10 bp) using FLASH (Magoč and Salzberg 2011)and length-filtered to retain only amplicons of the expected length ±2bp. After these filtering steps and read merging, the mean number of sequences retained per sample was 4109 (SD = 3032; median = 3526; range =135-29,863; counting only samples, positive and negative controls excluded). All remaining sequences were reduced to unique sequences using VSEARCH (Rognes et al. 2016).

Two thresholds for establishing successful detection and identification were considered based on the number of reads assigned to each cluster. First, the cluster had to have more sequences than any of the negative controls. This was not an issue because most negative controls were blank and only some had up to two sequences. Second, the cluster had to have more sequences than any possible mistag within each batch. To assess mistagging, we included all possible tag combinations which should not occurred within each batch in the demultiplexing step. This primarily includes the forward and reverse tag combinations which are not used within the batch because positive and negative controls have their own tags (Kitson et al. 2019), but also any combination of forward and reverse tags between the standard set of tags and the set of tags used for positive and negative controls. The level of mistagging detected in this way was between 78-436 sequences.

A single cluster from 701 samples (64%) passed this strict threshold. The clusters were then further clustered globally at 100% similarity to reduce the number of BLAST searches. Each sequence was then subjected to a BLAST search on a custom database collated from our own reference library sequences (included in supplementary data files as a spreadsheet (SequencesTable.csv in supplementary data files) and GenBank sequences of insects with emphasis on Diptera and Hymenoptera, common contaminants, and broad representatives of all kingdoms of life. Sequences from 649 samples were identified as one of the *Drosophila* species with more than 99% similarity. All positive controls passed the mistagging threshold and correctly identified the sample.

Based on reviewer comment we explored whether a less tight length filtering of sequences than ±2bp would change the outcome of the analysis as possibly too tight filtering could exclude parasitoid species with insertions or deletions in the amplicon sequence. Indeed, some chalcid parasitoids in our reference database have a 6bp deletion in the target locus. We repeated the analysis with less strict length filtering of ±6bp (which corresponds to ~±15% of amplicon length as our amplicon is 421bp). This modification had no effect on the downstream analysis because it only added 2.6% of sequences per sample on average and never resulted in a positive detection compared to background mistagging levels. We examined any cases where this approach would add more than 5% of sequences to the sample compared to the main analysis and most of these were bacterial sequences, only three were tentative parasitoid sequences (but not even close to positive detection compared to background mistagging levels).

**Classic molecular methods**

Samples other than those used for DNA metabarcoding were extracted using plate extractions following Ivanova et al. (2006).

For identification of the host species in “core” samples which were not successfully identified using DNA metabarcoding and in “additional samples” we used a multiplex PCR approach. We developed the multiplex PCRs based on host diversity revealed by the DNA metabarcoding as well as sampling of live iso-female lines at the study sites. The multiplex PCRs were based on the ITS2 and COI regions and included eleven *Drosophila* species (Table S2.1). In the case of eight host species, we designed primers based on the ITS2 locus using a common forward primer and a specific reverse primer. For the remaining three species, we designed primers based on the COI locus with specific forward and reverse primers. All multiplex PCRs were first optimized as simple PCRs and their specificity was tested out with a special focus on the most phylogenetically related species. Then we combined primers into multiplex PCRs according to the length of products in order to minimise number of reactions. Products of different length were detected using gel electrophoresis. In total, we used four multiplex PCRs (M1-M4) and one simple PCR (S1). In all cases, the thermocycling conditions were 94 °C for 5 min, following by 30 cycles of 94 °C for 40 s, 52 °C for 30 s, and 72 °C for 40 s, and final extension of 72 °C for 2 minutes.

| **PCR name** | ***Drosophila* species** | **Primer names and sequences (3'-5')** | **Gene** | **Product size (bp)** |
| --- | --- | --- | --- | --- |
| M1 | birchii | BIRCOI_50F: AGGAATAGTAGGAACATCTC | COI | 528 |
|  |  | BIRCOI_536R: TAATACAGGTAAAGATAAAAGAAG |  |  |
|  | rubida | UNIVERSAL_F: TGCTTGGACTACATATGG | ITS2 | 360 |
|  |  | RUB_R: AAACGACATGAAAGATACTAG |  |  |
|  | pseudoananassae | PSACOI_F407: GCAATTTTTTCGCTTCATC | COI | 208 |
|  |  | PSACOI_R574: AATTTCGATCAGTTAGTAATATG |  |  |
|  | pandora | UNIVERSAL_F: TGCTTGGACTACATATGG | ITS2 | 93 |
|  |  | PAN_R: ATACCATATGCATACTATAATG |  |  |
| M2 | bunnanda | UNIVERSAL_F: TGCTTGGACTACATATGG | ITS2 | 371 |
|  |  | BUN_R: ATTTACAATTTGTTAGCCATTAAC |  |  |
|  | pallidifrons | UNIVERSAL_F: TGCTTGGACTACATATGG | ITS2 | 181 |
|  |  | PAL_R: CTCATGCAACAGAGGTA |  |  |
| M3 | simulans | UNIVERSAL_F: TGCTTGGACTACATATGG | ITS2 | 313 |
|  |  | SIM_R: TCCATTTAACGAACCAAC |  |  |
|  | sulfurigaster | UNIVERSAL_F: TGCTTGGACTACATATGG | ITS2 | 184 |
|  |  | SUL_R: TTTCTCATGCAACAGAGTA |  |  |
| M4 | immigrans | UNIVERSAL_F: TGCTTGGACTACATATGG | ITS2 | 365 |
|  |  | IM_R2: GCTGTCTATTTTAACAATTTG |  |  |
|  | pseudotakahashii | UNIVERSAL_F: TGCTTGGACTACATATGG | ITS2 | 260 |
|  |  | PST_R2: ACAATTCTAGATAATATTTTATGC |  |  |
| S1 | bipectinata | BIP_AND_PSA_F: TTTTTATAATGATATGCATATGG | COI | 288 |
|  |  | BIP_R: TGAGGTTGTATAAAGCAATG |  |  |

**Table S2.1**. Multiplex PCRs to identify host species: primer sequences, primer combinations and length of PCR products.

We cross-checked consistency of host identification between DNA metabarcoding and multiplex on 40 samples previously identified by DNA metabarcoding. The results matched, but multiplex PCRs showed no result in 16 cases and were inconclusive within the sulfurigaster complex (*D. sulfurigaster* and *D. pallidifrons*).

When multiplex PCR failed or was inconclusive, we used Sanger sequencing to identify the host species. We developed Diptera specific primers based on the ITS2 region: one forward primer ITS2DipF106 (TGCTTGGACTACATATGG) with two reverse primers ITS2DipR240-1 (ATTTTTTATGCTAGACATTTCTC) and ITS2DipR240-2 (TTTTTATGCTAGACATTCCTC) with the final product of 280 bp. The reverse primers were mixed in the PCR in 1:1 ratio. The thermocycling condition were 94 °C for 5 min, following by 30 cycles of 94 °C for 40 s, 50 °C for 30 s, and 72 °C for 40 s, and final extension of 72 °C for 2 minutes. We used this approach also to confirm all *D. sulfurigaster* and *D. pallidifrons* species identifications from the multiplex PCR.

**Parasitoid detection and identification using PCR and Sanger sequencing**

We developed and tested custom parasitoid detection primers based on the 28S D2 region. We used insect sequences publicly available in GenBank to identify candidate regions with specific differences between Hymenoptera parasitoids and their Diptera hosts and then tested the primers using samples from a wide range of wasp and fly species (Figure S2.1). The resulting primers we developed are HymF74 (GCCCAGCACTGAATCC) and HymR238 (CAAGCAACCCGACTCTAA), with a resulting product of 164bp. We used the primers with annealing temperature of 55°C, but it safely identifies parasitiods from 48°C to 62°C. We used the same primers, but in combination with D1_F (ACCCGCTGAATTTAAGCATAT; (Harry et al. 1996); general 28S D1 primer downstream from HymF74) and 28SD1R (CAACTTTCCCTTACGGTACT; (Larsen 1992); general 28S D2 primer upstream from HymR238). This allowed us to obtain longer wasp sequences from parasitoid positive samples (~210bp; exact length depends on parasitoid species), which we then matched using BLAST to our reference library (SequencesTable.csv in supplementary data files). Sequences were identified as one of the parasitoid species with more than 99% similarity.

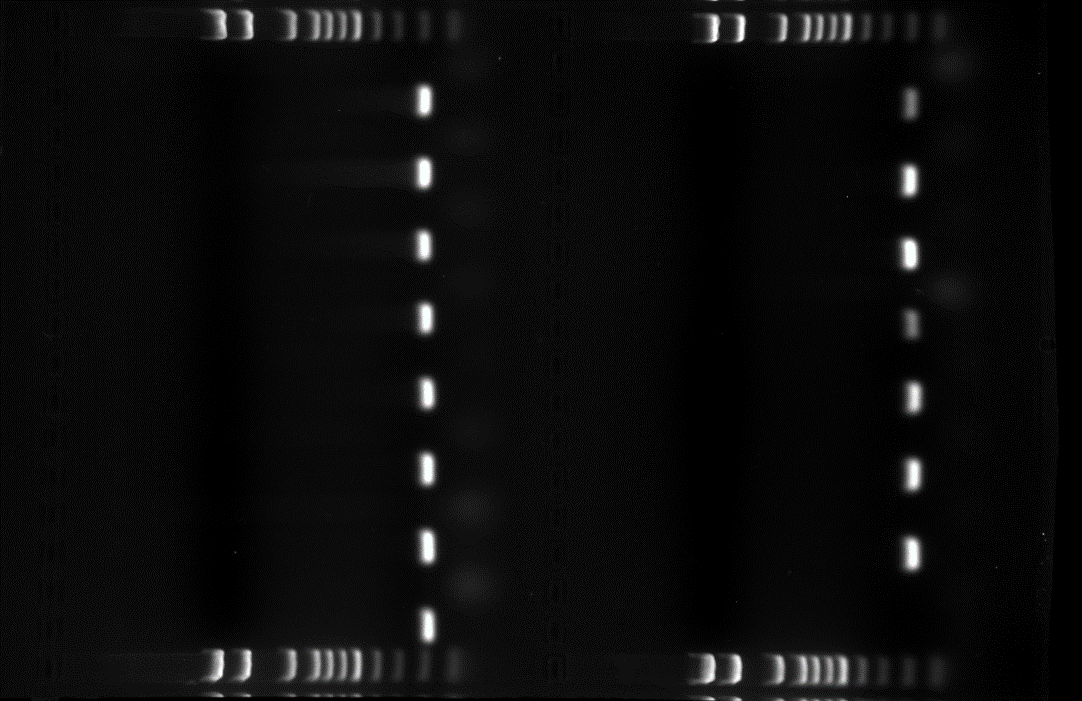

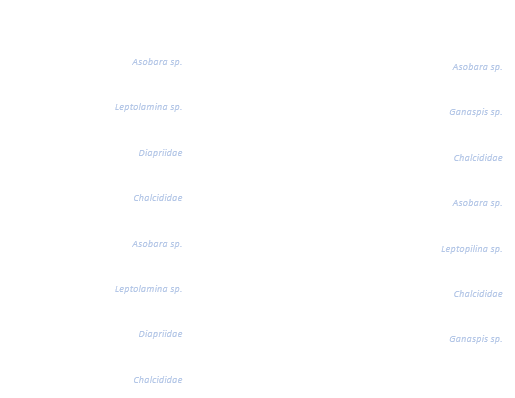

**Figure S2.1**. Test of the specificity of parasitoid detection primers. Photograph of 1.5% agarose gel showing PCR products (164 bp) obtained with HymF74 and HymR238 primers. Diptera (in white) and Hymenoptera (in blue) samples originated from Australian wet tropics. DNA ladder 100bp.

| **Parasitoid** | **Short Code** | **n** |
| --- | --- | --- |
| Asobara sp. 1 | A1 | 75 |
| Asobara sp. 2 | A2 | 21 |
| Asobara sp. 3 | A3 | 1 |
| Braconidae sp. 1 | B1 | 8 |
| Braconidae sp. 2 | B2 | 1 |
| Chalcidoidea sp. 1 | C1 | 3 |
| Diapriidae sp. 1 | D1 | 46 |
| Diapriidae sp. 2 | D2 | 5 |
| Diapriidae sp. 3 | D3 | 5 |
| Ganaspis sp. 1 | G1 | 11 |
| Leptolamina sp. 1 | LM1 | 83 |
| Leptopilina sp. 1 | LP1 | 146 |
| Leptopilina sp. 2 | LP2 | 10 |
| Leptopilina sp. 3 | LP3 | 5 |
| Pteromalidae sp. 1 | P1 | 16 |
| Unidentified | U | 7 |

**Table S2.2.** Parasitoid species identified, and the short codes used to annotate Figure 3 in the main text.

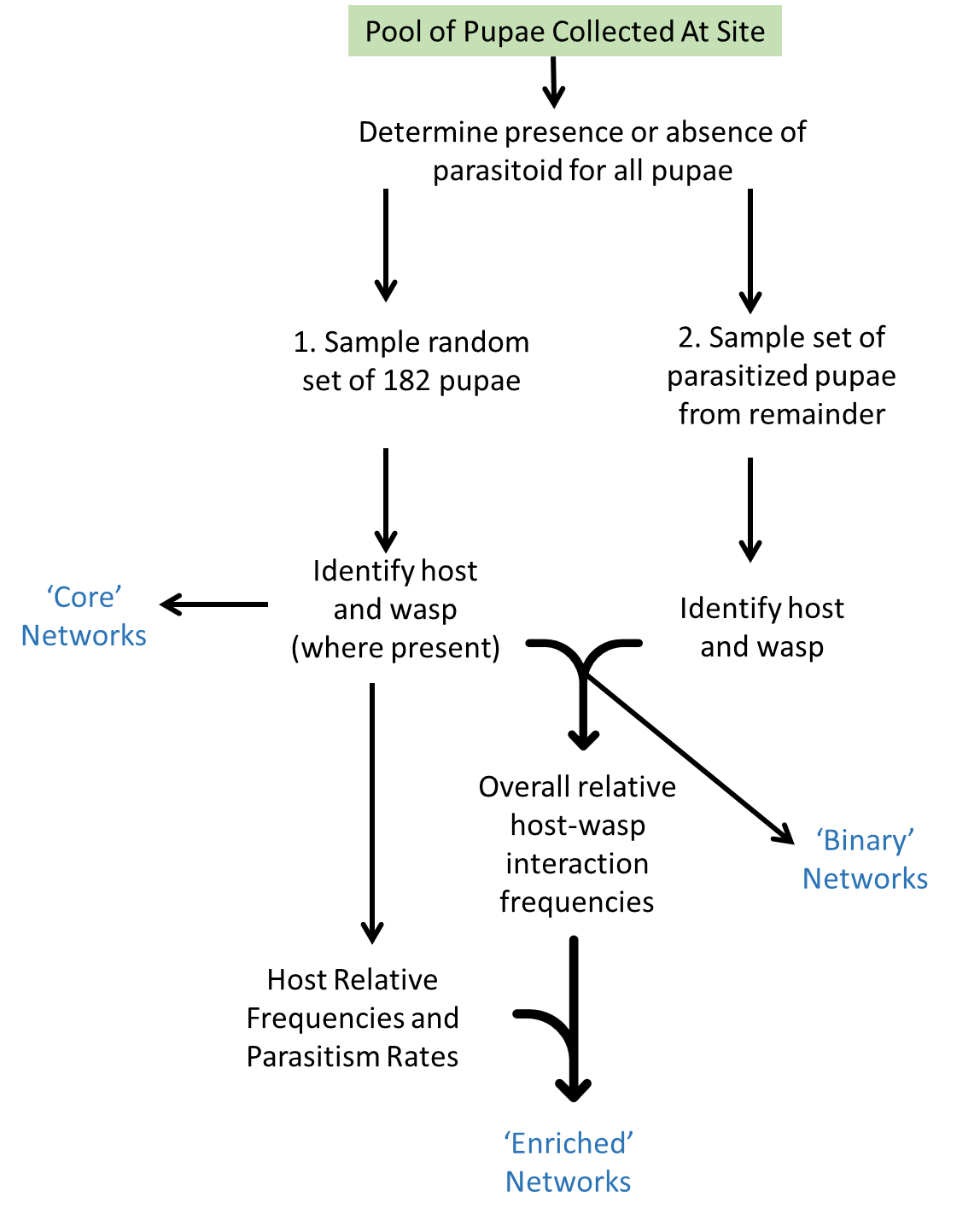

**Figure S2.2.** Flowchart of process to generate the interaction networks as described in the main text. The core networks were generated by random sampling of the available pool of pupae, and represent host frequencies and the parasitism rate on each species. Additional pupae, known to be parasitised were also sequenced to enrich the diversity of parasites observed attacking each host at each site.

Supplementary Information 3 - Statistical Analyses

### Introduction

This appendix contains additional data tables and the details of statistical analyses presented in the main text. Results are presented here as raw statistical outputs from R for maximum clarity. This is a dynamic document, generated using R Markdown. The source script for this file, as well as all material necessary to replicate the analysis are available at https://github.com/jcdterry/AusDrosTransect_CodeandData.

Raw identification data is available as SimplifiedFullData.csv, however note that due to our hierarchical sampling approach this should not be treated as a simple list of observed interactions. For long term archive and data-reuse purposes, tidied data and full explanatory metadata will also be made available in a standardised format through the Environmental Information Data Centre <https://doi.org/10.5285/85657c4c-54c9-4d02-a262-455ea1c38d95>.

### Impact of elevation on parasitism rates

#### Raw Data

Table S3.1. Raw counts of detailing parasitism frequency across all 14 sites.

| Site | Not-Parasitised | Parasitised | Total | Fraction Parasitised | Elevation | Transect |
| --- | --- | --- | --- | --- | --- | --- |
| K070 | 566 | 67 | 633 | 0.106 | 70 | K |
| K150 | 303 | 48 | 351 | 0.137 | 150 | K |
| K230 | 304 | 48 | 352 | 0.136 | 230 | K |
| K310 | 310 | 48 | 358 | 0.134 | 310 | K |
| K390 | 548 | 41 | 589 | 0.070 | 390 | K |
| K670 | 296 | 44 | 340 | 0.129 | 670 | K |
| K730 | 650 | 63 | 713 | 0.088 | 730 | K |
| P070 | 334 | 185 | 519 | 0.356 | 70 | P |
| P250 | 300 | 58 | 358 | 0.162 | 250 | P |
| P350 | 317 | 223 | 540 | 0.413 | 350 | P |
| P590 | 284 | 82 | 366 | 0.224 | 590 | P |
| P650 | 305 | 51 | 356 | 0.143 | 650 | P |
| P780 | 373 | 45 | 418 | 0.108 | 780 | P |
| P880 | 671 | 75 | 746 | 0.101 | 880 | P |

#### Statistical models - effect of transect and elevation

##### Model Selection

```
## Single term deletions
##
## Model:
## cbind(AllParasitismData$yes, AllParasitismData$no) ~ Transect *
##     Elevation
##                    Df Deviance F value Pr(>F)
## <none>                  125.15
## Transect:Elevation  1   158.15  2.6363 0.1355
```

Interaction term between transect and elevation non-significant and dropped from final model, but single terms retained.

```
## Single term deletions
##
## Model:
## cbind(AllParasitismData$yes, AllParasitismData$no) ~ Transect +
##     Elevation
##           Df Deviance F value   Pr(>F)
## <none>         158.14
## Transect   1   381.17  15.513 0.002317 **
## Elevation  1   304.22  10.160 0.008645 **
## ---
## Signif. codes:  0 '***' 0.001 '**' 0.01 '*' 0.05 '.' 0.1 ' ' 1
```

##### Final Model

| Model Term | Estimate | Std. Error |
| --- | --- | --- |
| (Intercept) | -1.5993 | 0.2639 |
| TransectP | 1.0578 | 0.2813 |
| Elevation | -0.0015 | 0.0005 |

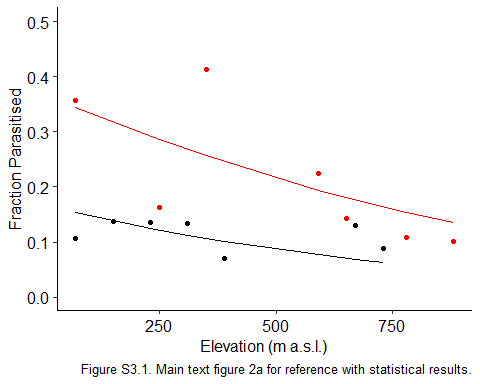

#### Incorporating host species ID

##### Data selection

For the within-species parasitism rare analysis we only include relatively common hosts: *D. pseudoananassae*, *D. rubida* and *D. pallidifrons*. Table S3.2 shows why these these species were selected - they were considerably more abundant and were the only three species found at all six sites in our survey.

Table S3.2. Abundance of fly species and presence across sites. Only three species are very widely distributed and are used in the species-specific parasitism rate analysis.

| Host | Number of Sites Present | Total |
| --- | --- | --- |
| immigrans | 1 | 5 |
| serrata | 1 | 1 |
| sulfurigaster complex | 1 | 1 |
| bunnanda | 2 | 6 |
| Drosophila sp. 1 | 2 | 6 |
| pseudotakahashii | 2 | 59 |
| pandora | 4 | 26 |
| bipectinata | 5 | 47 |
| birchii | 5 | 36 |
| sulfurigaster | 5 | 18 |
| pallidifrons | 6 | 133 |
| pseudoananassae | 6 | 222 |
| rubida | 6 | 486 |

Table S3.3. Site-specific parasitism data for three most abundant host species.

| Site | Host | Parasitised | Not-Parasitised | Total | Fraction Parasitised | Elevation | Transect |
| --- | --- | --- | --- | --- | --- | --- | --- |
| K070 | pallidifrons | 1 | 3 | 4 | 0.250 | 70 | K |
| K390 | pallidifrons | 4 | 16 | 20 | 0.200 | 390 | K |
| K730 | pallidifrons | 7 | 28 | 35 | 0.200 | 730 | K |
| P070 | pallidifrons | 4 | 6 | 10 | 0.400 | 70 | P |
| P350 | pallidifrons | 12 | 5 | 17 | 0.706 | 350 | P |
| P880 | pallidifrons | 9 | 38 | 47 | 0.191 | 880 | P |
| K070 | pseudoananassae | 7 | 25 | 32 | 0.219 | 70 | K |
| K390 | pseudoananassae | 9 | 54 | 63 | 0.143 | 390 | K |
| K730 | pseudoananassae | 4 | 23 | 27 | 0.148 | 730 | K |
| P070 | pseudoananassae | 80 | 10 | 90 | 0.889 | 70 | P |
| P350 | pseudoananassae | 0 | 8 | 8 | 0.000 | 350 | P |
| P880 | pseudoananassae | 0 | 2 | 2 | 0.000 | 880 | P |
| K070 | rubida | 34 | 45 | 79 | 0.430 | 70 | K |
| K390 | rubida | 15 | 39 | 54 | 0.278 | 390 | K |
| K730 | rubida | 28 | 38 | 66 | 0.424 | 730 | K |
| P070 | rubida | 9 | 6 | 15 | 0.600 | 70 | P |
| P350 | rubida | 165 | 57 | 222 | 0.743 | 350 | P |
| P880 | rubida | 23 | 27 | 50 | 0.460 | 880 | P |

##### Model Selection

Maximal model, with all two-way interactions between Host species, Elevation and Transect ID.

```
## Single term deletions
##
## Model:
## FracPara ~ Host + Transect + Elevation + Transect:Elevation +
##     Host:Elevation + Host:Transect
##                    Df Deviance F value Pr(>F)
## <none>                  33.250
## Transect:Elevation  1   47.083  3.3285 0.1055
## Host:Elevation      2   40.435  0.8645 0.4572
## Host:Transect       2   34.094  0.1016 0.9045
```

Sequentially dropping non-significant terms:

```
## Single term deletions
##
## Model:
## FracPara ~ Host + Transect + Elevation + Transect:Elevation +
##     Host:Elevation
##                    Df Deviance F value  Pr(>F)
## <none>                  34.094
## Transect:Elevation  1   55.033  6.1415 0.03264 *
## Host:Elevation      2   46.158  1.7691 0.21988
## ---
## Signif. codes:  0 '***' 0.001 '**' 0.01 '*' 0.05 '.' 0.1 ' ' 1
## Single term deletions
##
## Model:
## FracPara ~ Host + Transect + Elevation + Transect:Elevation
##                    Df Deviance F value  Pr(>F)
## <none>                  46.158
## Host                2   73.034  3.4937 0.06372 .
## Transect:Elevation  1   67.974  5.6717 0.03467 *
## ---
## Signif. codes:  0 '***' 0.001 '**' 0.01 '*' 0.05 '.' 0.1 ' ' 1
```

Transect:Elevation interaction term retained in minimal model:

```
## glm(formula = FracPara ~ Host + Transect + Elevation + Transect:Elevation,
##     family = "quasibinomial", data = CommonHostData, weights = Total)
```

| Model Term | Estimate | Std. Error |
| --- | --- | --- |
| (Intercept) | -1.4294 | 0.6115 |
| Hostpseudoananassae | 0.1722 | 0.5649 |
| Hostrubida | 0.9373 | 0.4660 |
| TransectP | 2.6537 | 0.5533 |
| Elevation | -0.0002 | 0.0009 |
| TransectP:Elevation | -0.0028 | 0.0012 |

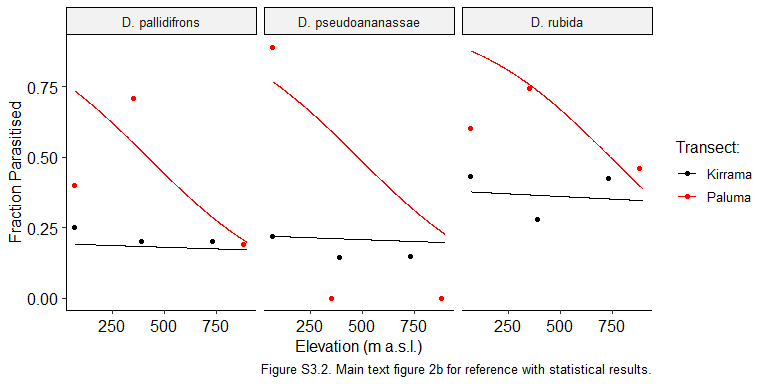

### Sampling completeness

Site-specific species accumulation curves show saturation of host species, but not for parasitoid wasps.

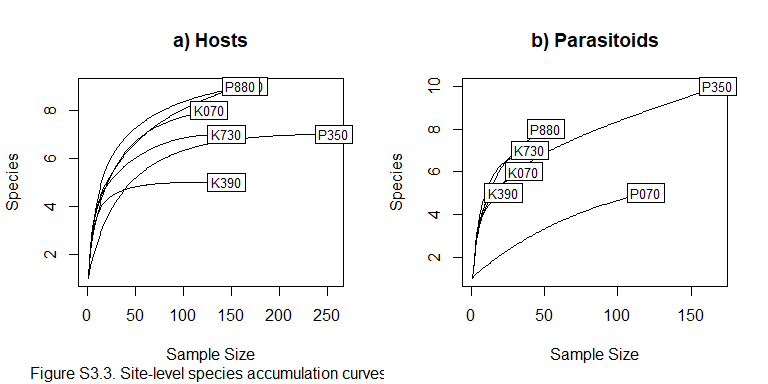

### Network metrics

#### Basic Specificity Statistics

##### Hosts

Table S3.4. Diversity of parasitoids that attack each host

| Host | n |
| --- | --- |
| bipectinata | 3 |
| birchii | 7 |
| bunnanda | 1 |
| Drosophila sp. 1 | 1 |
| immigrans | 1 |
| pallidifrons | 11 |
| pandora | 3 |
| pseudoananassae | 5 |
| pseudotakahashii | 7 |
| rubida | 11 |
| sulfurigaster | 3 |
| sulfurigaster complex | 1 |

##### Parasitoids

Table S3.5. Diversity of hosts that each parasitoid attacks

| LongName | n |
| --- | --- |
| Asobara sp. 1 | 2 |
| Asobara sp. 2 | 3 |
| Asobara sp. 3 | 1 |
| Braconidae sp. 1 | 1 |
| Braconidae sp. 2 | 1 |
| Chalcidoidea sp. 1 | 3 |
| Diapriidae sp. 1 | 8 |
| Diapriidae sp. 2 | 3 |
| Diapriidae sp. 3 | 3 |
| Ganaspis sp. 1 | 3 |
| Leptolamina sp. 1 | 6 |
| Leptopilina sp. 1 | 9 |
| Leptopilina sp. 2 | 3 |
| Leptopilina sp. 3 | 3 |
| Pteromalidae sp. 1 | 5 |

#### Plots

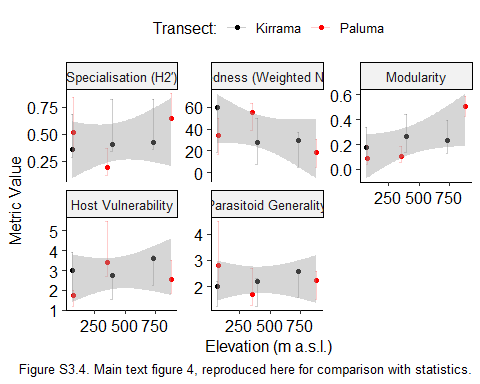

#### Statistical tests examining effect of elevation on network metrics

First testing with Transect (Kirrama or Paluma) as a co-predictor. Note the p-values reported here are unadjusted for multiple testing, so should be interpreted with due caution.

##### Raw Data

| Site | Elevation | Transect | generality.HL | H2 | Modularity | vulnerability.LL | weighted NODF |
| --- | --- | --- | --- | --- | --- | --- | --- |
| K070 | 70 | K | 2.000 | 0.358 | 0.176 | 3.000 | 59.524 |
| K390 | 390 | K | 2.200 | 0.407 | 0.266 | 2.750 | 28.125 |
| K730 | 730 | K | 2.571 | 0.425 | 0.234 | 3.600 | 29.383 |
| P070 | 70 | P | 2.800 | 0.513 | 0.089 | 1.750 | 34.409 |
| P350 | 350 | P | 1.700 | 0.192 | 0.102 | 3.400 | 55.490 |
| P880 | 880 | P | 2.250 | 0.650 | 0.504 | 2.571 | 18.640 |

### H2`

```
## lm(formula = Value ~ Elevation_km + Transect, data = filter(FiveMetricData,
##     Metric == "H2"))
```

| Model Term | Estimate | Std. Error | t-value | p-value |
| --- | --- | --- | --- | --- |
| (Intercept) | 0.3179 | 0.1367 | 2.3253 | 0.1026 |
| Elevation_km | 0.1984 | 0.2335 | 0.8495 | 0.4580 |
| TransectP | 0.0478 | 0.1425 | 0.3357 | 0.7592 |

##### Nestedness

```
## lm(formula = Value ~ Elevation_km + Transect, data = filter(FiveMetricData,
##     Metric == "weighted NODF"))
```

| Model Term | Estimate | Std. Error | t-value | p-value |
| --- | --- | --- | --- | --- |
| (Intercept) | 52.2992 | 11.9154 | 4.3892 | 0.0219 |
| Elevation_km | -33.5009 | 20.3514 | -1.6461 | 0.1983 |
| TransectP | -1.6024 | 12.4166 | -0.1291 | 0.9055 |

##### Modularity

```
## lm(formula = Value ~ Elevation_km + Transect, data = filter(FiveMetricData,
##     Metric == "Modularity"))
```

| Model Term | Estimate | Std. Error | t-value | p-value |
| --- | --- | --- | --- | --- |
| (Intercept) | 0.0806 | 0.0922 | 0.8745 | 0.4462 |
| Elevation_km | 0.3648 | 0.1575 | 2.3160 | 0.1035 |
| TransectP | -0.0069 | 0.0961 | -0.0719 | 0.9472 |

##### Vulnerability

```
## lm(formula = Value ~ Elevation_km + Transect, data = filter(FiveMetricData,
##     Metric == "vulnerability.LL"))
```

| Model Term | Estimate | Std. Error | t-value | p-value |
| --- | --- | --- | --- | --- |
| (Intercept) | 2.8093 | 0.5379 | 5.2224 | 0.0137 |
| Elevation_km | 0.7748 | 0.9188 | 0.8433 | 0.4610 |
| TransectP | -0.5713 | 0.5606 | -1.0191 | 0.3832 |

##### Generality

```
## lm(formula = Value ~ Elevation_km + Transect, data = filter(FiveMetricData,
##     Metric == "generality.HL"))
```

| Model Term | Estimate | Std. Error | t-value | p-value |
| --- | --- | --- | --- | --- |
| (Intercept) | 2.2321 | 0.3979 | 5.6094 | 0.0112 |
| Elevation_km | 0.0632 | 0.6796 | 0.0929 | 0.9318 |
| TransectP | -0.0095 | 0.4147 | -0.0228 | 0.9832 |

Repeating without transect co-predictor.

### H2`

```
lm( Value~Elevation_km,
    data = filter(FiveMetricData, Metric == 'H2') )%>% TidyOutput()
## lm(formula = Value ~ Elevation_km, data = filter(FiveMetricData,
##     Metric == "H2"))
```

| Model Term | Estimate | Std. Error | t-value | p-value |
| --- | --- | --- | --- | --- |
| (Intercept) | 0.3399 | 0.1059 | 3.2092 | 0.0326 |
| Elevation_km | 0.2031 | 0.2056 | 0.9876 | 0.3793 |

##### Nestedness

```
## lm(formula = Value ~ Elevation_km, data = filter(FiveMetricData,
##     Metric == "weighted NODF"))
```

| Model Term | Estimate | Std. Error | t-value | p-value |
| --- | --- | --- | --- | --- |
| (Intercept) | 51.5635 | 9.0865 | 5.6747 | 0.0048 |
| Elevation_km | -33.6587 | 17.6418 | -1.9079 | 0.1291 |

##### Modularity

```
## lm(formula = Value ~ Elevation_km, data = filter(FiveMetricData,
##     Metric == "Modularity"))
```

| Model Term | Estimate | Std. Error | t-value | p-value |
| --- | --- | --- | --- | --- |
| (Intercept) | 0.0775 | 0.0702 | 1.1037 | 0.3316 |
| Elevation_km | 0.3641 | 0.1363 | 2.6718 | 0.0557 |

##### Vulnerability

```
## lm(formula = Value ~ Elevation_km, data = filter(FiveMetricData,
##     Metric == "vulnerability.LL"))
```

| Model Term | Estimate | Std. Error | t-value | p-value |
| --- | --- | --- | --- | --- |
| (Intercept) | 2.5470 | 0.4746 | 5.3662 | 0.0058 |
| Elevation_km | 0.7186 | 0.9215 | 0.7798 | 0.4791 |

##### Generality

```
## lm(formula = Value ~ Elevation_km, data = filter(FiveMetricData,
##     Metric == "generality.HL"))
```

| Model Term | Estimate | Std. Error | t-value | p-value |
| --- | --- | --- | --- | --- |
| (Intercept) | 2.2277 | 0.3026 | 7.3611 | 0.0018 |
| Elevation_km | 0.0622 | 0.5876 | 0.1059 | 0.9207 |

Finally, testing again, this time including MatrixSize as a copredictor. This follows the method of Morris *et al.* (2004) cited in the main text, and is measured as the dimensions of the interaction matrix.

Note that the size of network was highly confounded by the transect:

| Site | Matrix_size |
| --- | --- |
| K070 | 24 |
| K390 | 20 |
| K730 | 28 |
| P070 | 35 |
| P350 | 40 |
| P880 | 40 |

### H2`

```
## lm(formula = Value ~ Elevation_km + Matrix_size, data = filter(MetricData_MatrixSize,
##     Metric == "H2"))
```

| Model Term | Estimate | Std. Error | t-value | p-value |
| --- | --- | --- | --- | --- |
| (Intercept) | 0.3274 | 0.3019 | 1.0842 | 0.3576 |
| Elevation_km | 0.1999 | 0.2473 | 0.8084 | 0.4780 |
| Matrix_size | 0.0004 | 0.0098 | 0.0454 | 0.9667 |

##### Nestedness

```
## lm(formula = Value ~ Elevation_km + Matrix_size, data = filter(MetricData_MatrixSize,
##     Metric == "weighted NODF"))
```

| Model Term | Estimate | Std. Error | t-value | p-value |
| --- | --- | --- | --- | --- |
| (Intercept) | 44.1383 | 25.4851 | 1.7319 | 0.1817 |
| Elevation_km | -35.5272 | 20.8753 | -1.7019 | 0.1873 |
| Matrix_size | 0.2631 | 0.8258 | 0.3186 | 0.7709 |

##### Modularity

```
## lm(formula = Value ~ Elevation_km + Matrix_size, data = filter(MetricData_MatrixSize,
##     Metric == "Modularity"))
```

| Model Term | Estimate | Std. Error | t-value | p-value |
| --- | --- | --- | --- | --- |
| (Intercept) | 0.1391 | 0.1964 | 0.7084 | 0.5298 |
| Elevation_km | 0.3796 | 0.1608 | 2.3603 | 0.0994 |
| Matrix_size | -0.0022 | 0.0064 | -0.3433 | 0.7540 |

##### Vulnerability

```
## lm(formula = Value ~ Elevation_km + Matrix_size, data = filter(MetricData_MatrixSize,
##     Metric == "vulnerability.LL"))
```

| Model Term | Estimate | Std. Error | t-value | p-value |
| --- | --- | --- | --- | --- |
| (Intercept) | 3.1482 | 1.2993 | 2.4230 | 0.0939 |
| Elevation_km | 0.8698 | 1.0643 | 0.8173 | 0.4736 |
| Matrix_size | -0.0213 | 0.0421 | -0.5060 | 0.6477 |

##### Generality

```
## lm(formula = Value ~ Elevation_km + Matrix_size, data = filter(MetricData_MatrixSize,
##     Metric == "generality.HL"))
```

| Model Term | Estimate | Std. Error | t-value | p-value |
| --- | --- | --- | --- | --- |
| (Intercept) | 2.3707 | 0.8583 | 2.7620 | 0.0700 |
| Elevation_km | 0.0982 | 0.7031 | 0.1397 | 0.8978 |
| Matrix_size | -0.0051 | 0.0278 | -0.1821 | 0.8671 |

### $\boldsymbol{\beta}$ - diversity

For the analysis of beta-diversity we used binary networks and Jaccard dissimilarity. Labeling of components of dissimilarity are described in the main text. To make use of additional information about host composition, separate from the interaction list, we use a customised version of bipartite::betalinkr(). This is detailed in ExtHostData_betalinkr.R, made available with all other code.

#### Multiple Regression on Distance Matrices

Communities do not differ more on different mountains or with greater elevational difference.

```
MRM(OS_Dist_Mat~Elevation_Dist+Mountain_Dff, nperm = 50000)$coef
##                 OS_Dist_Mat    pval
## Int            3.327439e-01 0.84670
## Elevation_Dist 4.958105e-05 0.59074
## Mountain_Dff   2.533025e-02 0.49316
MRM(WN_Dist_Mat~Elevation_Dist+Mountain_Dff, nperm = 50000)$coef
##                 WN_Dist_Mat    pval
## Int            0.7586218365 0.91722
## Elevation_Dist 0.0001404419 0.21048
## Mountain_Dff   0.0234319712 0.68846
MRM(ST_Dist_Mat~Elevation_Dist+Mountain_Dff, nperm = 50000)$coef
##                  ST_Dist_Mat    pval
## Int             4.258779e-01 0.67008
## Elevation_Dist  9.086082e-05 0.15728
## Mountain_Dff   -1.898283e-03 0.95200
MRM(SHost_Dist_Mat~Elevation_Dist+Mountain_Dff, nperm = 50000)$coef
##                SHost_Dist_Mat    pval
## Int              3.520339e-01 0.72150
## Elevation_Dist   8.925166e-05 0.45040
## Mountain_Dff    -1.312413e-02 0.78836
```

##### Repeating without transect difference as a check:

```
MRM(OS_Dist_Mat~Elevation_Dist, nperm = 50000)$coef
##                 OS_Dist_Mat    pval
## Int            3.538140e-01 0.70190
## Elevation_Dist 3.507056e-05 0.65496
MRM(WN_Dist_Mat~Elevation_Dist, nperm = 50000)$coef
##                 WN_Dist_Mat    pval
## Int            0.7781128825 0.74850
## Elevation_Dist 0.0001270188 0.22176
MRM(ST_Dist_Mat~Elevation_Dist, nperm = 50000)$coef
##                 ST_Dist_Mat    pval
## Int            4.242989e-01 0.51586
## Elevation_Dist 9.194825e-05 0.10822
MRM(SHost_Dist_Mat~Elevation_Dist, nperm = 50000)$coef
##                SHost_Dist_Mat    pval
## Int              3.411170e-01 0.83274
## Elevation_Dist   9.676985e-05 0.38182
```

##### Dissimilarity matrices plots

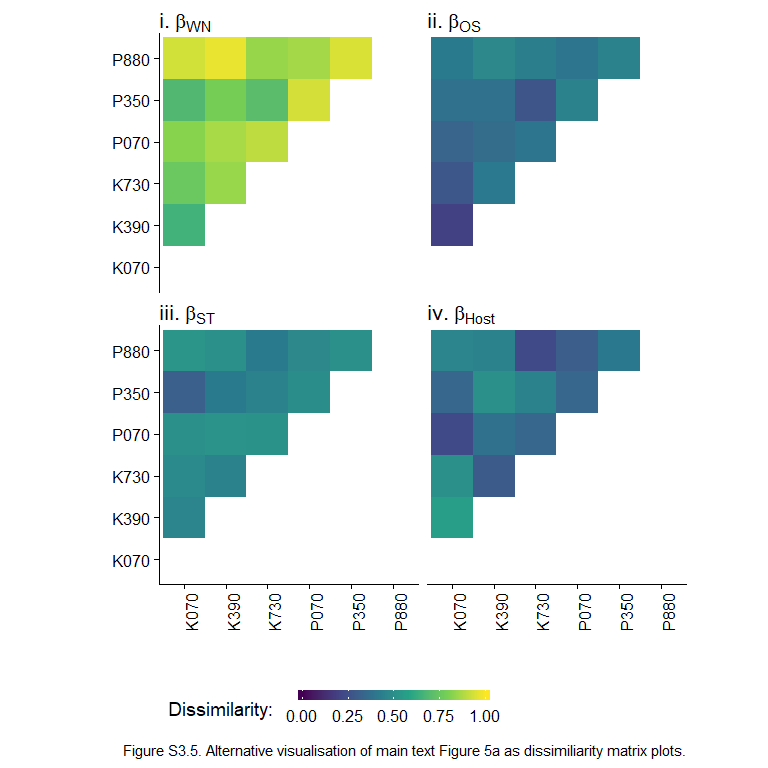

### Session Information

```
## R version 3.5.3 (2019-03-11)
## Platform: x86_64-w64-mingw32/x64 (64-bit)
## Running under: Windows 10 x64 (build 19041)
##
## Matrix products: default
##
## locale:
## [1] LC_COLLATE=English_United Kingdom.1252
## [2] LC_CTYPE=English_United Kingdom.1252
## [3] LC_MONETARY=English_United Kingdom.1252
## [4] LC_NUMERIC=C
## [5] LC_TIME=English_United Kingdom.1252
##
## attached base packages:
## [1] stats     graphics  grDevices utils     datasets  methods   base
##
## other attached packages:
##  [1] lme4_1.1-21          Matrix_1.2-15        broom_0.5.2
##  [4] cassandRa_0.1.0      ecodist_2.0.1        cowplot_0.9.4
##  [7] ggpubr_0.2           bipartiteD3_0.2.0    magrittr_1.5
## [10] knitr_1.23           readxl_1.3.1         bipartite_2.13
## [13] sna_2.4              network_1.15         statnet.common_4.3.0
## [16] vegan_2.5-5          lattice_0.20-38      permute_0.9-5
## [19] forcats_0.4.0        stringr_1.4.0        dplyr_0.8.1
## [22] purrr_0.3.2          readr_1.3.1          tidyr_0.8.3
## [25] tibble_2.1.3         ggplot2_3.2.0        tidyverse_1.2.1
## [28] RevoUtils_11.0.3     RevoUtilsMath_11.0.0
##
## loaded via a namespace (and not attached):
##  [1] httr_1.4.0         maps_3.3.0         viridisLite_0.3.0
##  [4] jsonlite_1.6       splines_3.5.3      dotCall64_1.0-0
##  [7] modelr_0.1.4       assertthat_0.2.1   highr_0.8
## [10] cellranger_1.1.0   yaml_2.2.0         pillar_1.4.1
## [13] backports_1.1.4    glue_1.3.1         downloader_0.4
## [16] digest_0.6.19      RColorBrewer_1.1-2 minqa_1.2.4
## [19] rvest_0.3.4        colorspace_1.4-1   htmltools_0.3.6
## [22] pkgconfig_2.0.2    haven_2.1.0        scales_1.0.0
## [25] mgcv_1.8-27        generics_0.0.2     withr_2.1.2
## [28] lazyeval_0.2.2     cli_1.1.0          crayon_1.3.4
## [31] evaluate_0.14      nlme_3.1-137       MASS_7.3-51.1
## [34] xml2_1.2.0         tools_3.5.3        hms_0.4.2
## [37] munsell_0.5.0      cluster_2.0.7-1    compiler_3.5.3
## [40] rlang_0.4.0        nloptr_1.2.1       grid_3.5.3
## [43] rstudioapi_0.10    spam_2.2-2         igraph_1.2.4
## [46] labeling_0.3       rmarkdown_1.13     boot_1.3-20
## [49] gtable_0.3.0       R6_2.4.0           lubridate_1.7.4
## [52] stringi_1.4.3      parallel_3.5.3     Rcpp_1.0.1
## [55] fields_9.8-3       tidyselect_0.2.5   xfun_0.8
## [58] coda_0.19-2
```
